# Deuterium Magnetic Resonance Spectroscopy of Early Treatment-Induced Changes in Tumour Lactate *in vitro*

**DOI:** 10.1101/2021.04.03.438324

**Authors:** Josephine L Tan, Daniel Djayakarsana, Hanzhi Wang, Rachel W Chan, Colleen Bailey, Angus Z Lau

## Abstract

Elevated production of lactate is a key characteristic of aberrant tumour cell metabolism and can be non-invasively measured as an early marker of tumour response using deuterium (^2^H) magnetic resonance spectroscopy (MRS). Following treatment, changes in the ^2^H-labeled lactate signal could identify tumour cell death or impaired metabolic function, which precede morphological changes conventionally used to assess tumour response. In this work, the association between apoptotic cell death, extracellular lactate concentration, and early treatment-induced changes in the ^2^H-labeled lactate signal was established in an *in vitro* tumour model. Experiments were conducted at 7 T on acute myeloid leukemia cells which had been treated with 10 µg/mL of the chemotherapeutic agent cisplatin. At 24 and 48 hours after cisplatin treatment, the cells were injected with 20 mM of [6,6’-^2^H_2_]glucose and scanned over two hours using a two-dimensional ^2^H MR spectroscopic imaging sequence. The resulting signals from ^2^H-labeled glucose, lactate, and water were quantified using a spectral fitting algorithm implemented on the OXford Spectroscopy Analysis (OXSA) MATLAB toolbox. After scanning, the cells were processed for histological stains (TUNEL [terminal deoxynucleotidyl transferase UTP nick end labeling] and H&E [hematoxylin and eosin]) to assess apoptotic area fraction and cell morphology respectively, while a colorimetric assay was used to measure extracellular lactate concentrations in the supernatant. Significantly lower levels of ^2^H-labeled lactate were observed in the 48-hour treated cells compared to the untreated and 24-hour treated cells, and these changes were significantly correlated with an increase in apoptotic fraction and a decrease in extracellular lactate. By establishing the biological processes associated with treatment-induced changes in the ^2^H-labeled lactate signal, these findings suggest that ^2^H MRS of lactate may be valuable in evaluating early tumour response.

## Introduction

Imaging plays a growing role in the non-invasive assessment of tumour response to treatment. For most solid tumours, the accepted clinical standard for response evaluation is based on the measurement of lesion size by computed tomography (CT) and magnetic resonance imaging (MRI) ^1,2^. However, morphological changes may not occur until several weeks to months after treatment and do not always accurately reflect tumour progression or regression, rendering it difficult to assess early tumour response ^3,4^. Imaging techniques that can detect early-stage markers of tumour response are therefore of great interest for optimizing subsequent therapy and reducing the burden of ineffective or unnecessary treatments ^5^.

A potential early-stage imaging marker of treatment response is lactate ^6–8^. In contrast to normal cells, tumour cells exhibit aberrant metabolism characterized by an increase in glycolysis regardless of aerobic conditions ^9^, resulting in elevated lactate concentrations as high as 30 mM in multiple tumour types ^8^. Following treatment, changes in lactate levels can provide insight on biological processes associated with response that precede gross changes in tumour size, including tumour cell death or impairment of metabolic function ^10^.

Recently, deuterium (^2^H) magnetic resonance spectroscopy (MRS) was introduced as a potential clinical tool for imaging metabolism *in vivo* following the non-invasive administration of deuterated substrates such as glucose ^11–13^. Compared to fluorine-18 fluorodeoxyglucose (^18^F-FDG) positron emission tomography (PET), ^2^H MRS can be performed on the same system as anatomical MRI scans, is non-radioactive, and informs on metabolism beyond glucose uptake. In addition, the short relaxation times of ^2^H-labeled molecules and low natural abundance of the ^2^H nucleus contribute to the well-resolved resonances that can be achieved in the ^2^H NMR spectra ^11–14^. By detecting downstream ^2^H-labeled metabolites like lactate, ^2^H MRS has been shown to differentiate between normal and glioblastoma tissue in humans ^12^ and detect treatment response 48 hours post-chemotherapy in murine tumour models ^13^. While these findings suggest that ^2^H MRS of lactate may be valuable for assessing tumour response to therapy, more work is needed to establish the biological processes associated with this metabolic change.

In this work, *in vitro* experiments were performed using ^2^H MRS to assess any correlations between ^2^H-MRS measures of lactate signal, extracellular lactate concentrations, and apoptotic cell death following chemotherapy. We show that significant decreases in the ^2^H-lactate signal can be observed at 48 hours after treatment and were associated with apoptotic cell death and decreased extracellular lactate. These results provide insight into the use of ^2^H MRS to assess tumour response at an earlier stage in therapy.

## Methods

### Cell preparation

The experimental workflow is summarized in Figure 1. An acute myeloid leukemia (AML-5) cell line was chosen for this *in vitro* study because of its well-documented apoptotic nature of response to cisplatin ^15–19^. AML cells were grown in suspension in twelve flasks containing 150 mL alpha minimum essential medium (Invitrogen Canada Inc., Burlington, Canada), with 5% fetal bovine serum (Fisher Scientific, Ottawa, Canada), and 1% penicillin and streptomycin (Invitrogen Canada Inc.). Flasks were maintained at 37°C and 5% CO_2_ until they reached confluence (∼10^6^ cells/mL). To induce apoptosis, cells in each of the flasks were treated with 10 µg/mL cisplatin 24 or 48 hours prior to imaging. A single sample was prepared by combining four flask volumes (∼6 x 10^8^ cells total), which were centrifuged at 3000 rpm for 15 minutes at 4°C using a fixed angle centrifuge (Beckman Coulter, Brea, CA, USA). After removal of excess supernatant, the pellet was resuspended in phosphate-buffered saline (PBS) and centrifuged at 2400g for 10 min at 4°C using a swinging bucket centrifuge (Thermo Electron Corp., Asheville, NC, USA). PBS was used to resuspend the pellet into an MR-compatible glass tube with a final volume of 600 µL. Each experiment utilized three samples. 10 minutes prior to the ^2^H scan, 100 µL of 140 mM [6,6’-^2^H_2_]glucose solution (Cambridge Isotope Laboratories, Tewksbury, MA, USA) was injected into each tube for a final ^2^H-labeled glucose concentration of 20 mM (and final volume=700 µL) and micropipetted 10 times in the solution to ensure mixing throughout the tube. The experiment was repeated twice to determine reproducibility and was performed with untreated samples and samples treated with cisplatin 24 and 48 hours prior to scanning.

**Figure 1:**
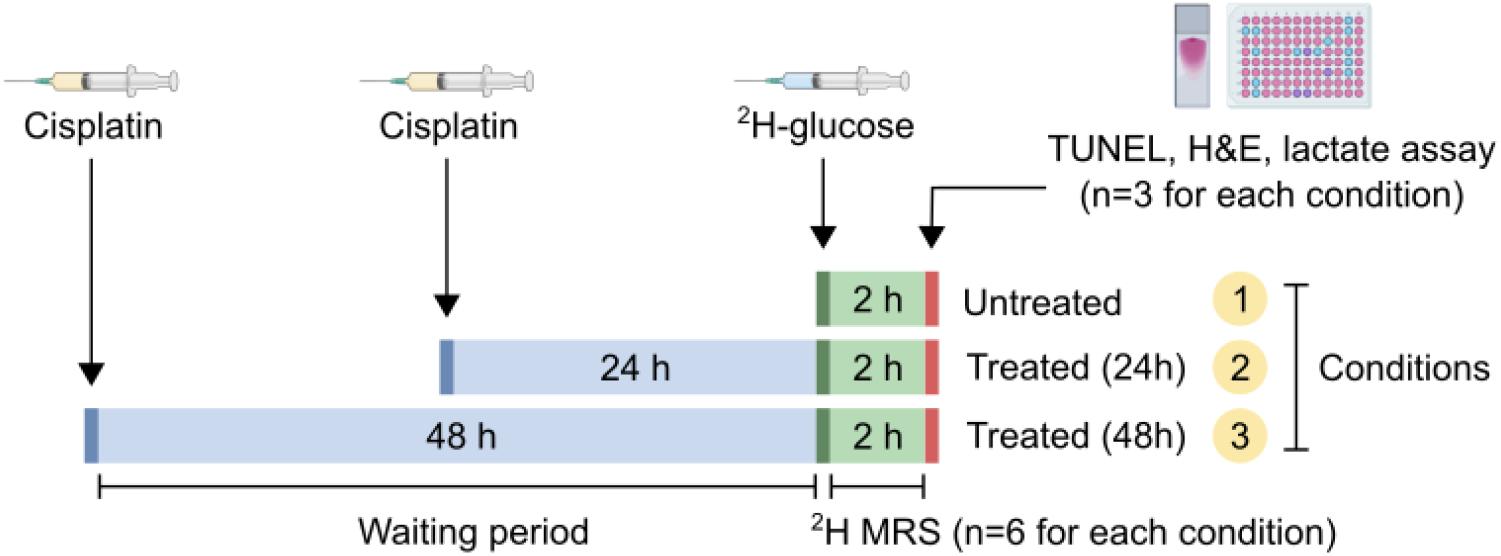
Schematic experimental timeline for the untreated and cisplatin-treated cells.

### ^2^H MRS data acquisition

MR experiments were performed at 7 T (Bruker BioSpec 7 T). A home-built transmit/receive surface coil (46.1 MHz, diameter 20 mm) was used for acquiring the ^2^H signal. The coil included a proton blocking trap circuit to prevent the induction of currents on the ^2^H coil by the ^1^H body coil in the scanner. Three MR-compatible glass tubes (6 mm diameter × 50 mm length) containing the cell samples were placed side-by-side in the center of the surface coil, as illustrated in Figure 2A. Magnetic field homogeneity was optimized with shim localized around the centre of the tubes. ^2^H signal excitation was achieved with a 90° rectangular radiofrequency pulse followed by 2D phase-encoding gradients (0.5 ms). The flip angle was calibrated using separate phantom scans prior to *in vitro* scans. 2D ^2^H MRS images were acquired at an in-plane resolution of 10×3 mm^2^ and temporal resolution of 2 min (FOV=8×3 cm^2^, spectral width 3 kHz, 768 spectral points, flip angle=90°, TR=300 ms, 300 averages). The rectangular voxels were chosen to match the length of the cell sample tubes. A single ^2^H MRS image was first acquired to measure baseline metabolism before the cell samples were removed from the scanner and injected with deuterated glucose. Following the addition of glucose, the cell sample tubes were placed back onto the coil and reshimmed. 10 minutes after the injection of glucose, dynamic ^2^H MRS images were acquired over 120 mins. Proton T_1_-weighted images at an in-plane resolution of 0.03125×0.03125 cm^2^ (FOV=8×8 cm^2^, TR=51 ms, TE=2 ms) were acquired with the ^1^H volume coil.

**Figure 2:**
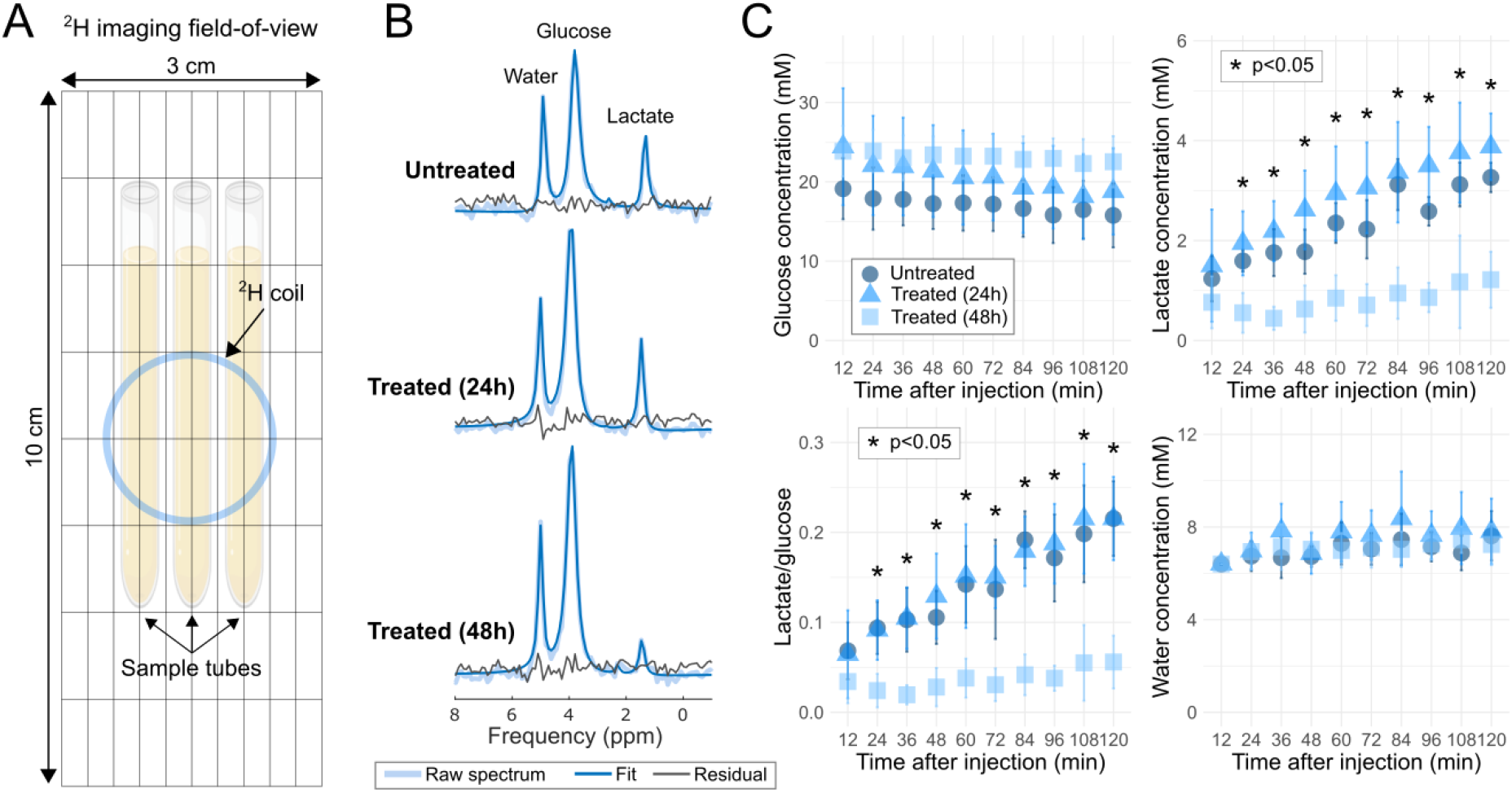
**(A)** Experimental setup consisting of three AML sample tubes lying on top of the ^2^H surface coil (blue circle), with overlaid 2D matrix. **(B)** Representative single voxel ^2^H spectra for the untreated and cisplatin-treated samples two hours after injection of [6,6’-^2^H_2_]glucose. Spectra are averaged over 12 minutes of acquisition. **(C)** Time course of 2H-labelled glucose, lactate, lactate/glucose and water concentrations 10 minutes after injection of [6,6’-^2^H_2_]glucose. Error bars are the standard deviation across 6 samples. Stars indicate a statistically significance difference between the 48-hour treated and untreated samples, and between the 48-hour and 24-hour treated samples (p<0.05).

### ^2^H MR data analysis

The spectrum for each sample tube was chosen from the voxel in the most sensitive region of the surface coil with the highest SNR. Spectra were averaged from every 12 min of data acquisition (corresponding to 30 images) to maximize SNR. Spectral fitting was performed using the ‘Advanced method for accurate, robust and efficient spectral fitting’ (AMARES) algorithm implemented in the OXford Spectroscopy Analysis (OXSA) MATLAB toolbox ^20^. The fitting model is constrained by prior knowledge including the relative chemical shifts (bounded by ±0.2 ppm) and linewidths. A Lorentzian line shape was fitted to each peak. Metabolite signals were quantified by area integration of the peaks, which included deuterium-labeled glucose (3.7 ppm), glutamate and glutamine (Glx, 2.4 ppm), lactate (1.4 ppm), and water (4.7 ppm). The fitting was bound between ±0.1 ppm. These chemical shifts were taken from literature ^11–13^ and verified from our spectral fitting. For display, ^2^H MRS maps were resampled and overlaid onto the corresponding proton T_1_-weighted image.

Signals were corrected by a factor that accounted for partial signal saturation, given by

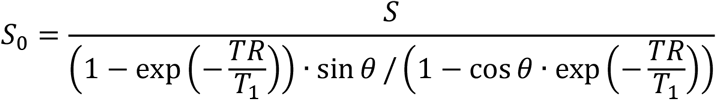

where S and S_0_ is the signal before and after correction, respectively, TR is the repetition time [ms], T_1_ is the longitudinal relaxation [ms], θ is the prescribed flip angle [rad], and TE is the echo time [ms]. The T_1_ of ^2^H-labeled metabolites were based on *in vivo* values from literature measured at 7 T ^12^: 64 ms (glucose), 146 ms (Glx), 297 ms (lactate), and 320 ms (water). Concentrations of labeled glucose and lactate were quantified by normalizing the integrals of the fitted peaks to that of the baseline water signal, which has a natural ^2^H abundance of 12.8 mM. This value is based on the 55.5 M concentration of water, the two protons of the water molecule, and 0.0115% natural abundance of deuterium. As described by de Graaf et al. ^12^, the absolute concentrations of glucose, Glx, lactate, and water were determined by normalizing to the average number of deuterons per molecule (glucose 2, Glx 1.33, lactate 3, water 2). Tukey’s post hoc test was used to determine statistically significant differences (p<0.05) in metabolite concentrations between the untreated and cisplatin-treated samples at every time point (12, 24, 36, 48, 60, 72, 84, 96, 108, 120 minutes) following glucose injection. Statistical calculations were performed with R (v4.0.3: R Core Team (2020), Vienna, Austria).

### *In vitro* assays

^2^H MRS data were compared to cell sample histology and lactate assay measurements. Immediately following ^2^H MRS, 3 samples from each treatment category (untreated, 24-hour treated, and 48-hour treated) we re transferred into 1.5 mL Eppendorf tubes and centrifuged at 2400g for 10 mins. 300 µL of supernatant was extracted from each tube and stored at -80°C for at least 48 hours for lactate assays. To prepare samples for histology, pellets were fixed in 10% formalin for at least 96 hours. The pellets were embedded in 3% agarose gel and stained with terminal deoxynucleotidyl transferase-mediated 2’-deoxyuridine 5’-triphosphate nick-end labeling (TUNEL) and hematoxylin and eosin (H&E) to assess apoptotic area and cell morphology, respectively (University Health Network Pathology Research Program, Toronto, Canada). Slides were examined under a light microscope (10x and 200x magnification). ImageJ2 ^21^ was used to measure the percent area of TUNEL positive (TUNEL+) staining in three fields of view of each slide.

The supernatant lactate concentration was measured using the Lactate Colorimetric Assay Kit (Biovision Inc., Milpitas, CA, USA, #K607) according to the manufacturer’s instructions. For sample preparation, 0.8 µL of the supernatant was added per well. Tukey’s post hoc test was used to determine statistically significant differences (p<0.05) between the untreated and cisplatin-treated samples for ^2^H-labeled lactate concentrations at the last MRI time point (120 minutes), and corresponding TUNEL+ staining and supernatant lactate concentrations.

## Results

The ^2^H spectra acquired two hours after the injection of [6,6’-^2^H_2_]glucose in the untreated and cisplatin-treated samples are presented in Figure 2B. The spectra were averaged from a 12-minute window. Shimming resulted in a linewidth of 15 to 20 Hz (0.050 to 0.070 ppm). Signals were observed from deuterated water (4.7 ppm), the injected glucose (3.7 ppm) and downstream lactate (1.4 ppm), but no Glx was detected in either the cisplatin-treated or untreated groups. Figure 2C shows the concentration of deuterated water and the ^2^H-labeled metabolites over two hours 10 minutes after ^2^H-glucose administration, averaged across 6 samples. The 48-hour treated sample exhibited reduced glucose metabolism, as indicated by steady concentrations of glucose and lactate over time. In contrast, glucose levels decreased while lactate levels increased over the two-hour time course of the 24-hour treated and untreated samples. Across 6 samples, a significantly lower lactate concentration was detected at all time points beyond 12 minutes (up to 120 min) after glucose administration in the 48-hour treated cells compared to those untreated (p<0.05) and compared to the 24-hour treated cells (p<0.05). Similar trends were observed in the time course of lactate/glucose ratios, where the ratios were significantly lower in the 48-hour treated cells compared to those untreated (p<0.05) and compared to the 24-hour treated cells (p<0.05) at all time points beyond 12 minutes post glucose injection. No significant differences in lactate or lactate/glucose were detected between the untreated and 24-hour treated samples at any time point (p>0.05). There were also no significant differences detected in glucose or water concentration at any time point between any of the samples. Figure 3 shows representative 2D spectroscopic images of metabolite signals two hours after glucose administration. Reduced lactate signal was observed in the 48-hour treated samples compared to the untreated and 24-hour treated samples.

**Figure 3:**
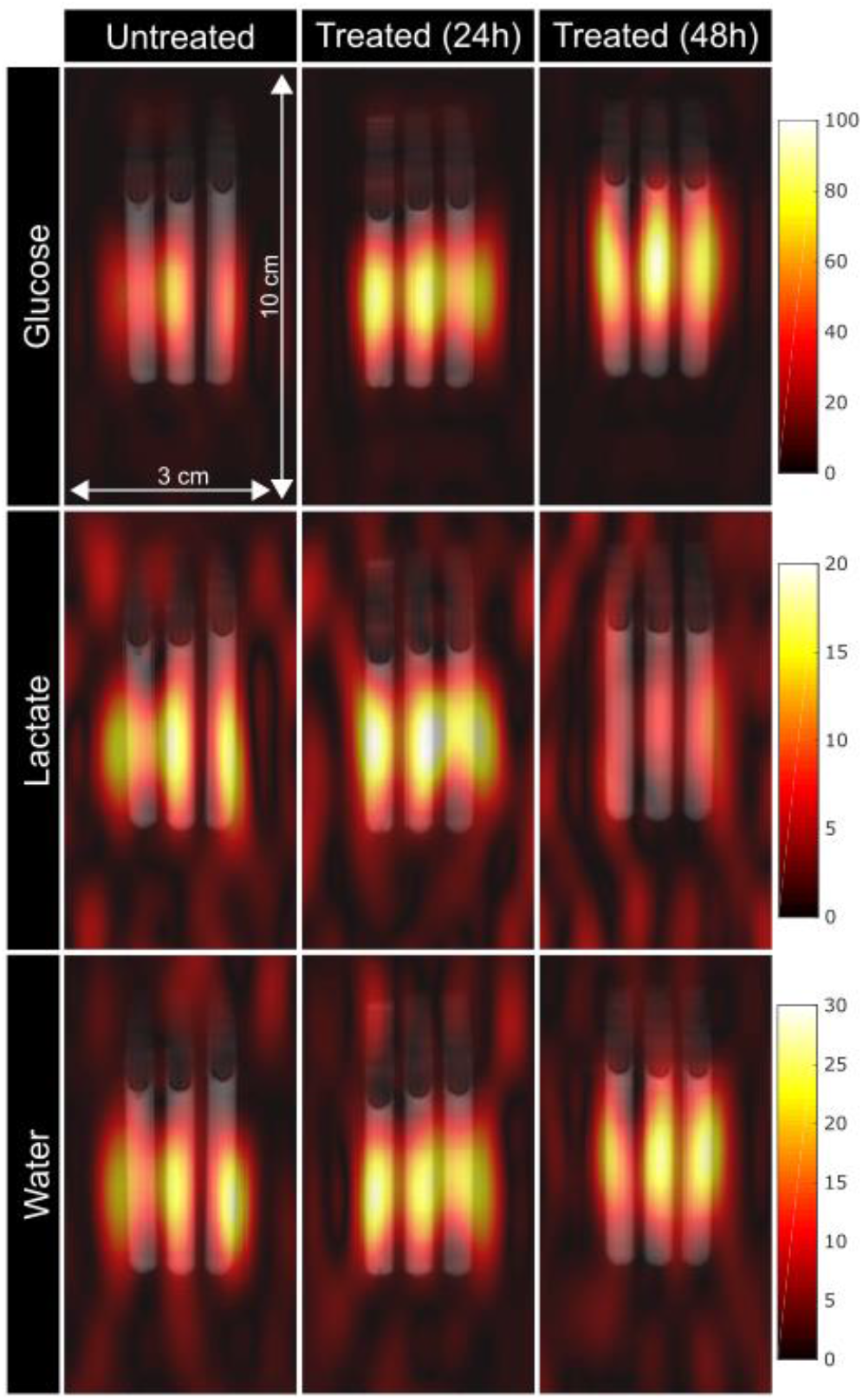
Representative ^2^H spectroscopic images overlaid on T_1_ images two hours after injection of [6,6’-^2^H_2_]glucose. The spectral data was averaged over 12 minutes of acquisition. The untreated, 24-hour treated, and 48-hour treated spectroscopic images shown were acquired from three separate experiments. Colourbars represent arbitrary signal intensity.

Figure 4 shows a microscope image of AML cells in suspension, which are approximately 10 µm in diameter. Apoptotic changes are evident 48 hours after treatment with cisplatin, such as nuclear condensation, membrane blebbing, and cell size shrinkage. These characteristics are reflected in the histology slides shown in Figure 5. Cell size shrinkage may have been caused by formalin fixation. The H&E sections of the 48-hour treated cells showed increased nuclear fragmentation, evidenced by dark purple clusters, compared to the untreated and 24-hour treated cells. Similarly, the TUNEL sections of the 48-hour treated cells showed increased positive staining, which appear as brown regions, compared to the untreated and 24-hour treated cells. Figure 6 compares the lactate concentrations measured by ^2^H MRS two hours after glucose injection from 3 samples, and corresponding lactate measurements from the assay and TUNEL+ area fraction from the histology slides. Across the 3 samples two hours after glucose injection, ^2^H-labeled lactate concentration was significantly lower in the 48-hour treated sample (1.6±0.4 mM) compared to the untreated (3.2±0.4 mM, p<0.05) and 24-hour treated samples (4.0±0.6 mM, p<0.05). There was no significant difference detected in ^2^H MRS lactate measurements between the untreated and 24-hour treated samples (p>0.05). The lactate assay of the sample supernatants indicated that there was less extracellular lactate in the 48-hour treated sample (7.0±2.1 mM) compared to untreated (11.3±1.0 mM) and 24-hour treated samples (11.5±2.3 mM). However, the differences were not significant between any group (p>0.05). In contrast, there was a significantly greater TUNEL+ area in the 48-hour treated cell pellet histology slides (16.8±4.6 %) compared to the untreated (3.9±0.8 %, p<0.05) and 24-hour treated slides (2.3±0.8 %, p<0.05). Like the ^2^H MRS lactate measurements, there was no significant difference in TUNEL+ area between the untreated and 24-hour treated samples (p>0.05). Figure 7 shows the correlation plots between ^2^H MRS lactate measurements, lactate assay measurements, and TUNEL+ area fraction. The ^2^H MRS lactate levels were significantly correlated with the assay measurements (p<0.05). Both MRS and assay lactate data were significantly negatively correlated with the TUNEL+ area fraction (p<0.05).

**Figure 4:**
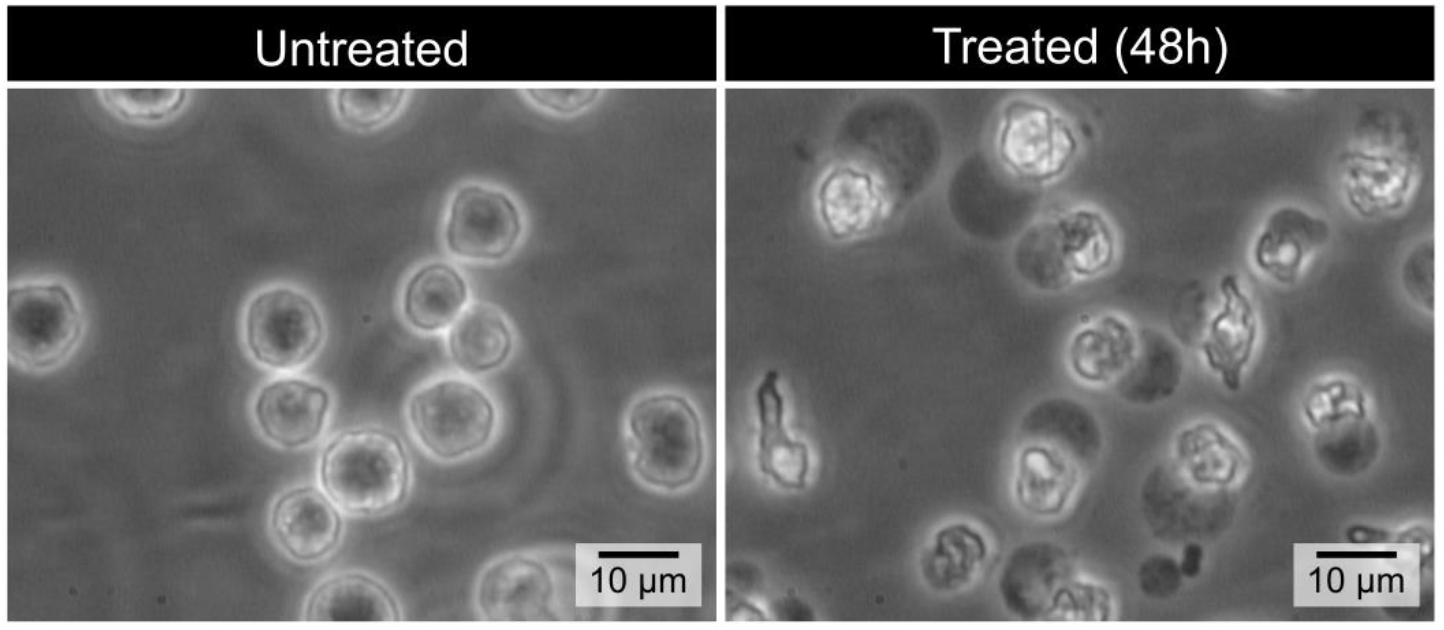
Phase contrast microscope images of AML cells in suspension. Cells are approximately 10 µm in diameter. Apoptotic changes can be observed at 48 hours after treatment including nuclear condensation, membrane blebbing, and cell size shrinkage.

**Figure 5:**
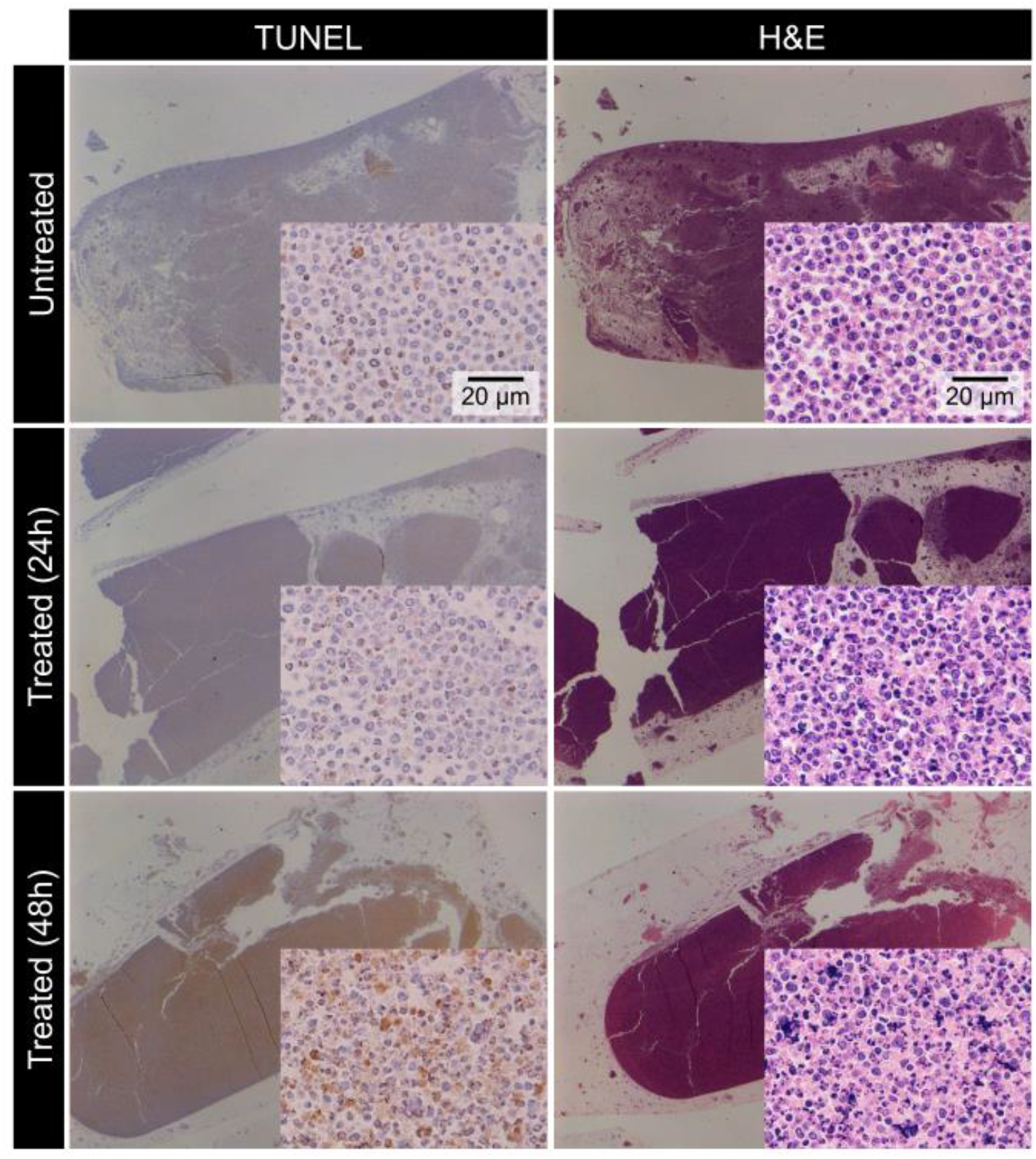
Representative TUNEL and H&E stained sections (10x and 200x magnification) from untreated and cisplatin-treated cell pellets, which were fixed in formalin for at least 96 hours after scanning. Formalin fixation may have caused cell shrinkage.

**Figure 6:**
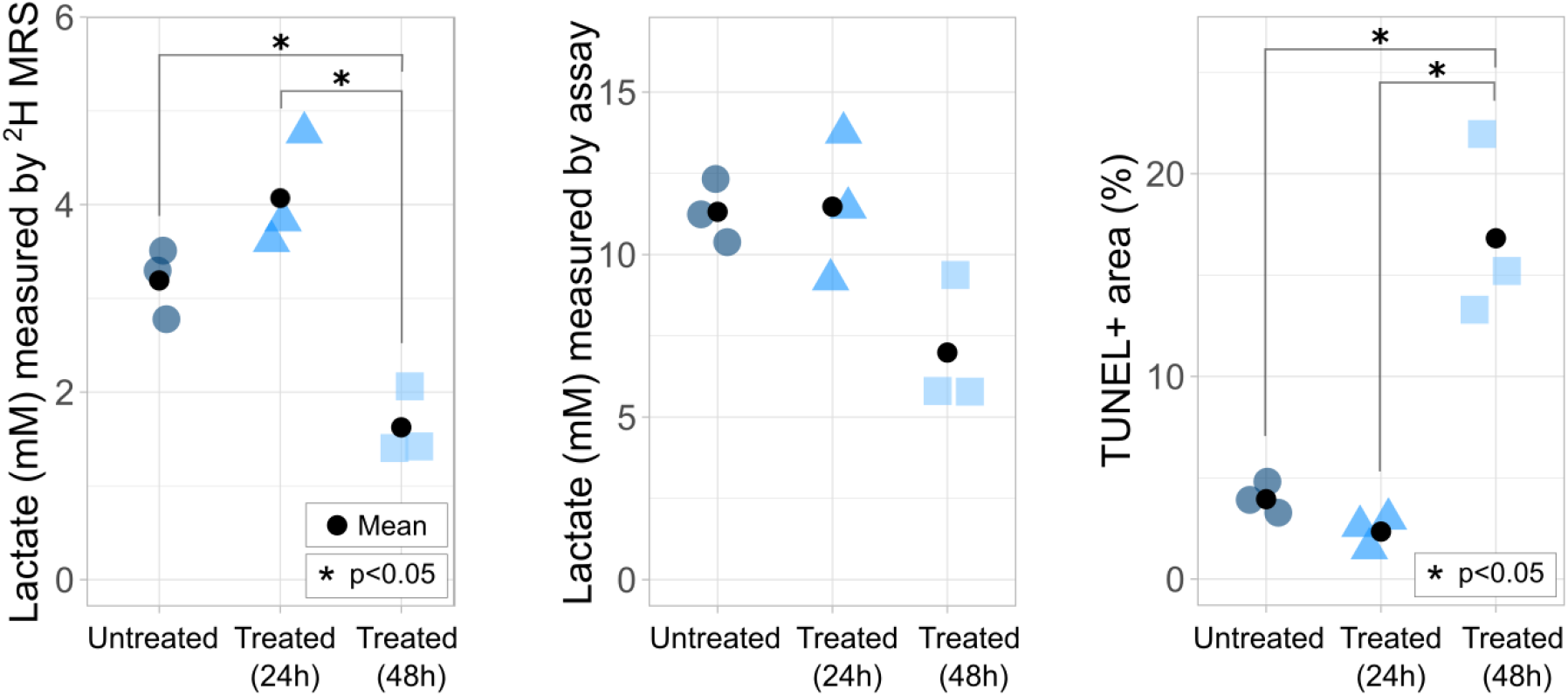
Comparison of ^2^H-lactate measured by ^2^H MRS 120 minutes after glucose injection, and corresponding assay measurements of lactate in supernatant and TUNEL positive percent area across 3 samples. ^2^H-lactate data is averaged from 12 minutes of data acquisition. Statistically significant differences (p<0.05) were determined using Tukey’s post hoc test.

**Figure 7:**
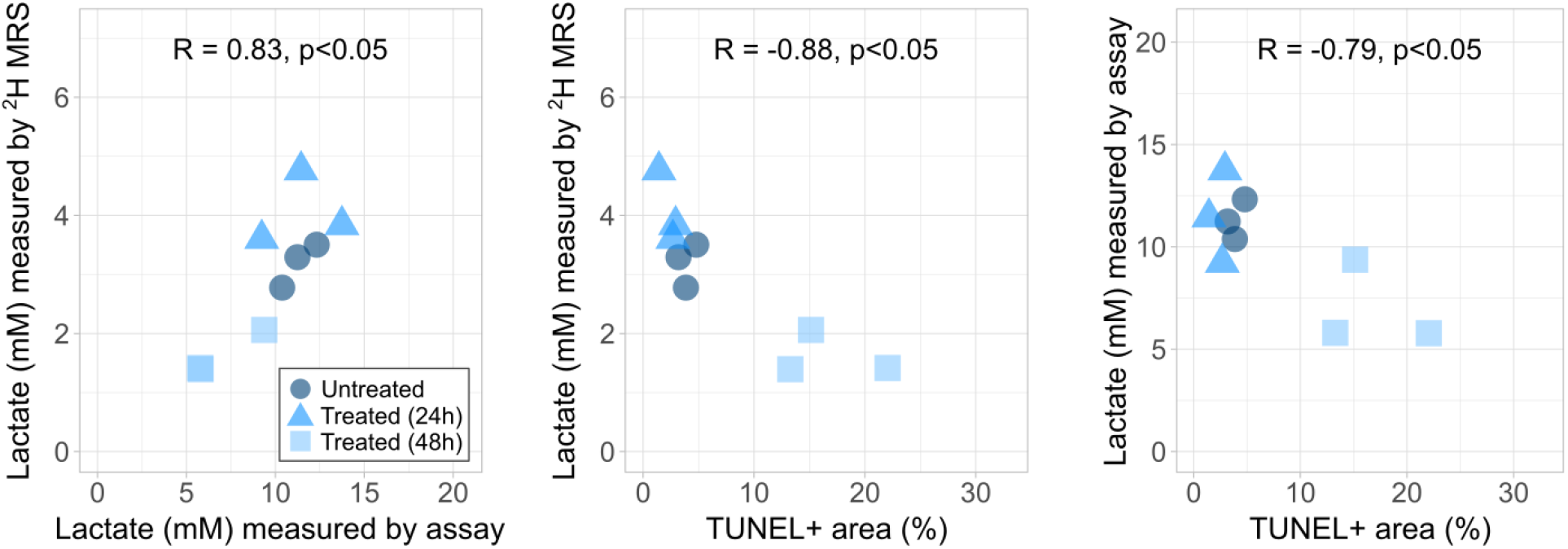
Significant correlations were determined between the lactate MRS and assay measurements, the MRS measurements and TUNEL positive percent area (TUNEL+ area), and the assay measurements and TUNEL+ area (p<0.05). R represents Pearson’s correlation coefficient.

## Discussion

In this study, the metabolic changes in cisplatin-treated AML cells were measured using ^2^H MRS and correlated with biological parameters including extracellular lactate concentration and apoptotic area fraction. Consistent with the Warburg effect ^9^ and previous ^2^H MRS studies of glucose metabolism in tumours ^12,13^, the AML cells demonstrated metabolic conversion of glucose into lactate without detectable amounts of Glx produced. This conversion was observed within two hours following the administration of ^2^H-labelled glucose, in agreement with other dynamic ^2^H MRS studies ^11,13^. Beyond 12 minutes after glucose administration, there was a significant decrease in ^2^H-labeled lactate production in the samples treated with cisplatin 48 hours prior to scanning, relative to the untreated samples and 24-hour treated samples. No significant differences in glucose concentrations were observed at any time point between any of the samples, although glucose utilization appeared reduced in 48-hour treated cells compared to the untreated and 24-hour treated cells. To account for the potential variation in ^2^H-glucose dosage and/or viable cell population within the sample, labeled lactate/glucose over time was also examined. The lactate/glucose ratio was significantly lower in the 48-hour treated samples compared to both the untreated and 24-hour treated samples beyond 12 minutes post-injection of labeled glucose. At 120 minutes post-injection of labeled-glucose, the lactate/glucose ratio across 6 samples for the untreated cells (0.22±0.04) and 48-hour treated cells (0.06±0.03) were lower than ^2^H-labeled lactate/glucose ratios measured in EL4 tumour bearing mice 48 hours after treatment with etoposide (0.27±0.12 to 0.12±0.06) ^13^. Reasons for this discrepancy might include the difference between cell lines, chemotherapeutic agents and doses, and variations arising from *in vivo* versus *in vitro* measurements such as oxygen supply and availability of glucose.

To validate the lactate concentrations quantified by ^2^H MRS, a colorimetric assay was used to measure lactate in the sample supernatant. In agreement with the ^2^H MRS data, lactate concentrations were lower in the supernatant of the 48-hour treated sample compared to the untreated and 24-hour treated samples. While there were no significant differences in assay-based lactate concentrations between any group, these measurements were significantly correlated to those measured by ^2^H MRS two hours after glucose administration. The assay-based lactate concentrations were an average of 3.6 times higher than the ^2^H MRS measurements, which may be explained by the presence of unlabeled lactate in the supernatant – for example, produced from residual cell medium that contains unlabeled glucose – which is undetected by ^2^H MRS. This effect should be considered when interpreting the ^2^H lactate signal in ^2^H MRS studies.

The decrease in lactate concentration, as measured by both ^2^H MRS and the lactate assay, was significantly correlated with the histologically confirmed increase in percent area of TUNEL positive staining 48 hours after cisplatin was administered. The 48-hour treated samples also demonstrated morphological features of apoptosis in H&E-stained sections. This finding suggests that an increase in cisplatin-induced apoptosis results in a decrease in lactate production since there are fewer viable cells to metabolize the administered glucose. However, given the percent of viable cells in the untreated (∼95%) and 48-hour treated (∼85%) samples (approximated from the TUNEL positive percent area), the change in lactate concentration between these two groups is greater than expected (∼0.3 mM [expected] vs. 1.6 mM [measured by ^2^H MRS] and 4.3 mM [measured by assay]). The observed drop in lactate concentration may be attributed to other cisplatin-mediated mechanisms in addition to apoptosis, such as a “stun” in glucose metabolism from disrupted cell machinery. Cisplatin, a chemotherapeutic drug known to exert cytotoxicity by apoptosis, has also been postulated to act as an anti-metabolic agent. Cisplatin was found to suppress the expression of glycolysis-related proteins including glucose transporters 1 and 4 (GLUT1 and GLUT4) and lactate dehydrogenase B (LDHB), leading to a decline in glucose uptake and lactate production *in vitro* ^22^. There may also be reduced expressions of other glycolysis-related proteins such as monocarboxylate transporters (MCT) and lactate dehydrogenase A (LDHA), which facilitate the transport of lactate into and out of the cell and the bidirectional conversion of pyruvate to lactate, respectively ^8^. Expressions of such proteins have been found to be significantly reduced in tumours treated with radiation compared to controls ^23^. Cell metabolism may also be arrested once apoptosis is initiated. For example, two glycolysis-limiting enzymes, phosphofructokinase and pyruvate kinase, have been shown to be inhibited by caspases during apoptosis ^24^. These events may affect lactate production before TUNEL staining can indicate apoptotic cell death. Western blot analysis of these protein levels in future ^2^H MRS studies may add valuable information regarding the mechanisms of the observed metabolic change.

The setup of this experiment consisted of a plane of three tubes placed horizontally above the centre of a ^2^H surface coil. This layout enabled sufficient signal acquisition using a 2D CSI sequence. Sensitivity was the greatest in the centre of the coil, leading to a high signal in a single voxel from each tube. For this reason - and to prevent potential signal contamination from neighbouring voxels - only those three voxels were chosen for analysis. In our case, 2D maps obtained via spectroscopic imaging served to demonstrate the spatial resolution and localization of the ^2^H signal relative to the coil within the three identical sample tubes. In a clinical application, these metabolic maps have been shown to distinguish between normal and aberrant metabolism with striking contrast ^12^. Using 3D ^2^H MRS maps of glucose, lactate, and Glx, de Graaf et al. have demonstrated elevated lactate and reduced Glx in a glioma relative to normal brain tissue ^12^. Given the nominal voxel size of 2×2×2 mm^3^ used in this study, these ^2^H MRS metabolic maps could be used to provide a quantitative assessment of response across the tumour volume and capture the presence of spatial metabolic heterogeneities that may further inform on tumour status. Due to its ability to map metabolites downstream of glucose, ^2^H MRS overcomes a major disadvantage of ^18^F-FDG PET which is limited to imaging glucose uptake ^10^.

These *in vitro* experiments were not impacted by factors such as tumour vascularity which introduce additional complexity for metabolite quantification ^11,13^. However, while the *in vitro* approach permitted isolation of the cellular effects of cisplatin treatment, the results of this experiment may not be reflective of *in vivo* results. For example, the experiment was performed at room temperature instead of at the physiologically relevant temperature of 37°C. Accordingly, lactate production rates have been shown to vary between *in vivo* and *in vitro* environments ^25^. The glucose administered is also four times that found in normal blood, which may affect metabolic rates relative to those found in physiological conditions. In addition, our technique was tested on a single AML cell line which is highly glycolytic ^26^ and, being cancer of the blood, is not assessed for response via anatomical changes as done in solid tumours ^27^. These differences should be considered if comparing AML treatment response to solid tumours or cells that are less glycolytic. However, similar metabolic trends to this study were observed with ^2^H MRS in a murine lymphoma (EL4) model ^13^, in a glioma model (RG2), and in patients with glioblastoma multiforme (GBM) ^12^. Other limitations for this study include the varying number of cell passages. At a higher passage number, the cells may experience genetic drift that alters metabolism due to changes in morphology, growth rates, and protein expression ^28^. Despite the potential passage effects, the MR and biological data show consistent metabolic trends across the untreated, 24-hour treated, and 48-hour treated groups.

## Conclusions

Increased lactate levels are a consequence of aberrant metabolism exhibited by many tumour types that contribute to the survival and metastatic potential of tumour cells. The results of this work showed that significant treatment-induced changes in lactate can be detected by ^2^H MRS as early as 48 hours after chemotherapy and were associated with changes in extracellular lactate levels and apoptotic cell death. These findings suggest that ^2^H-labeled lactate may be used as an early marker of tumour response. Future work is needed to determine the mechanisms of the observed metabolic change.

## Acknowledgements

The authors gratefully acknowledge funding from NSERC (RGPIN-2017-06596). AML-5 cell line was kindly provided by Dr. Gregory Czarnota (Sunnybrook Health Sciences Centre).

